# Evaluating peer-to-peer bioinformatics education: a case study of student learning outcomes and community impact in an undergraduate multi-omic data analysis course

**DOI:** 10.1101/2025.10.07.680870

**Authors:** Wade R. Boohar, Kayla Y. Xu, Nicole Black, Mahija Mogalipuvvu, Kate Manley, Peter Calabrese, Jerry S.H. Lee

**Author notes:** Corresponding authors (WRB), (KYX). These authors contributed equally to this work (Wade R. Boohar, Kayla Y. Xu).

## Abstract

Computational methodology has become ubiquitous in biomedical research with the rise of big data analysis and popularity of artificial intelligence and machine learning. However, undergraduate bioinformatics education has largely struggled to keep pace with the demand for bioinformatics skills, due to a combination of social and resource-based barriers. In this case study, we discuss the application and outcomes of Multi-Omic Data Analysis, a peer-to-peer learning-based undergraduate bioinformatics course offered by the Department of Quantitative and Computational Biology at the University of Southern California. Over eight semesters, a cohort of student instructors taught 2-3 weekly lectures to 107 undergraduate students in the Quantitative Biology Bachelor of Science degree program. Lectures covered a range of topics, including R and Python data analysis, scientific communication, and general research readiness as undergraduate students. We find that bioinformatics education courses structured around peer-to-peer learning have great potential to overcome many of the obstacles to comprehensive undergraduate bioinformatics education, and provide additional benefits related to student cohesion and community. We further discuss the longevity and feasibility of such courses, both specific to our program and in undergraduate universities at large.

**Author Summary:** There is a growing need for accessible and beginner-friendly bioinformatics education at the undergraduate level. Traditional coursework often fails to meet this demand due to disciplinary separation between biology and computer science, as well as limited institutional resources. We evaluate a unique approach involving a student-led undergraduate bioinformatics course at the University of Southern California to understand how peer-to-peer learning influences student development in computational biology.

We examined over 100 students across eight academic semesters. Students varied in prior coding experience and research exposure. Through survey responses, we found that students left the course feeling more confident and prepared for research in faculty labs. Students reported improved competence in bioinformatics skills compared to before they took the course. We found that a major reason for this success was the peer-to-peer learning format. Students mentioned that learning from fellow students made the material more approachable and created a supportive community where they felt comfortable asking questions. Our work demonstrates that student-led initiatives can be a highly effective solution for filling critical gaps in university curricula, making bioinformatics education more accessible.

## Introduction

In the past decade, biomedical research has seen a significant shift towards computational analysis. With the rise of big data and increasing computational power, many research institutions have established dedicated departments to bioinformatics research, and bioinformatics in the private sector is expected to nearly triple its market size by 2030 [1]. As such, the accessibility of comprehensive bioinformatics education is now a necessity at the undergraduate level [2].

Such education, however, has not matched the growth of bioinformatics utility. In 2022, only 104 of 3,976 United States Title IV degree-granting institutions offered bachelor’s degrees in “Biomathematics, Bioinformatics, and Computational Biology” [3]. Of the 3,082 total degrees awarded in such categories, only 21.7% were bachelor’s degrees, illustrating how the vast amount of computational biology and bioinformatics education is still constrained to graduate-level programs. This contrasts the more ubiquitous degrees in biology, of which 136,148 bachelor’s degrees were awarded in 2022, and degrees in computer sciences, of which 112,408 bachelor’s degrees were awarded in the same timeframe (see supporting information for data and definition of degrees).

Traditional divisions of standard undergraduate degrees separate biological and computational sciences. Even universities which offer bioinformatics majors often require students to take separate classes in the computer science department that lack biological context or application. Students are then left to their own devices in learning how to apply computer science principles to biological problems. This can cause biology students to view computational work as being too difficult or foreign, resulting in poor learning outcomes [4].

One solution for addressing this problem is through peer-to-peer (PTP) learning strategies. PTP learning is a teaching tool that emphasizes interpersonal learning between peers, whether that be coworkers or undergraduate students. This approach has already been embraced in areas such as medical education [5]. One study found that undergraduate nursing students who were randomly assigned to a PTP model performed significantly better than their expert-led counterparts [6]. Other studies have found that undergraduate engineering students who serve as mentors in PTP programs also benefit as well [7]. Despite the mounting evidence in support of PTP-based programs in undergraduate curriculum, there is still little interest paid to programs which are fully taught by undergraduate students. One such program, DeCal at the University of California, Berkeley, does exist, and allows undergraduate students to propose and implement their own course plan for other students. However, to our knowledge, no bioinformatics course exists within DeCal.

Benefits of a PTP-based model for bioinformatics education can be multi-faceted. For universities with limited funding, employing students as educators allows for implementation without drawing resources from other classes. Motivation for participation in instruction can be given in the form of research credit, fulfilling a requirement that is common in many higher learning institutions for STEM fields [8, 9]. Current students may also serve as less intimidating mentor figures to underclassmen, allowing for increased interpersonal engagement that can serve as a vital tool for both learning and student retention.

Studies also show that PTP learning can improve retention of marginalized students within STEM by mitigating the impact of imposter syndrome, a feeling of anxiety caused by a disconnect between perceived internal success and external results [10]. Many computational courses have a reputation for being intellectually intimidating, especially if students have no prior experience programming. PTP learning can reduce this perception by allowing students to work with student instructors who have recently taken the courses themselves, allowing them to share strategies that helped them succeed. This unique viewpoint may improve overall enthusiasm and confidence by lowering the risks of mistakes or poor performance. Students who are falling behind in the class may also be more likely to reach out to instructors due to the shared undergraduate experience.

PTP classes also allow for more comprehensive and beginner-friendly introductions to bioinformatics. Such beginner-level courses can build a solid foundation in basic necessary skills and knowledge, mitigating the need for remedial support in later, more technical courses. PTP-based learning, with instructors who can have different research experiences, increases the breadth of skills that students are exposed to.

This article is a case study in one implementation of a current PTP-learning course at the University of Southern California (USC). Over a period of eight semesters, a student-led class known as Multi-Omic Data Analysis (MODA) was taught to 107 undergraduate students in the Quantitative Biology (QBIO) major using publicly available cancer data from The Cancer Genome Atlas [11]. We show that this PTP class improved students’ preparedness for and perception of bioinformatics research through the teaching of foundational scientific methodologies and hands-on work with genomic, transcriptomic, epigenomic, and proteomic data.

## Results

A total of 107 students were involved in the program over the course of 8 semesters (Fig 1a). 83 of those students responded to a pre-semester survey, 70 responded to a post-semester survey, and 13 responded to the follow-up alumni survey. Most students took the course in their second or third year (Fig 1b). Many students enrolled in MODA after switching into the QBIO degree program partway through their undergraduate education. For these students, this class may have been their first exposure to bioinformatics.

**Fig 1.**
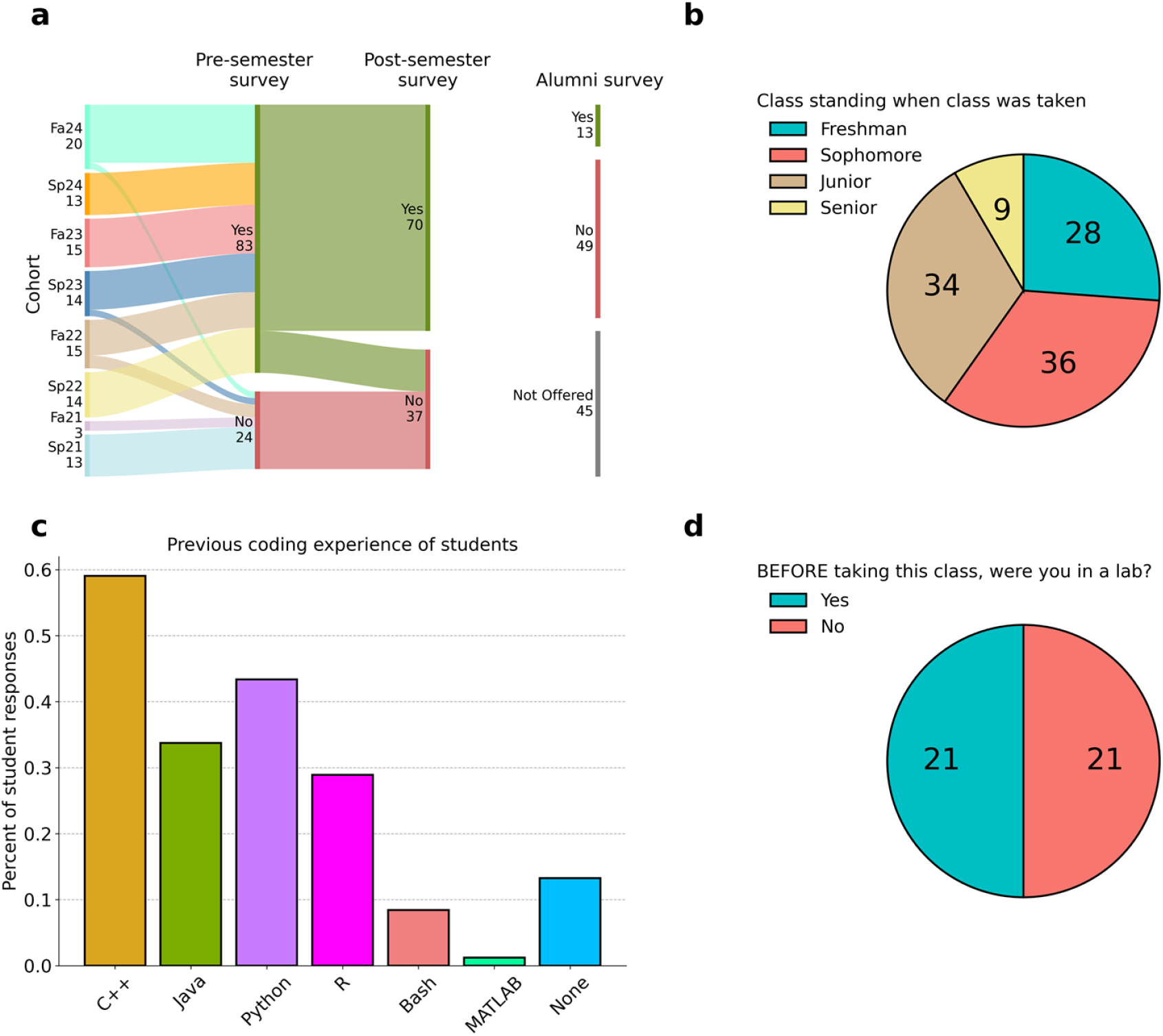
Demographics of students who participated in the pre- and post-semester survey for the student-led MODA section. (a) Sankey plot of all students grouped by cohort and whether they responded to the pre- and/or post-semester surveys. Alumni survey responses were anonymous, and as such, no cohort information was collected. Survey questions were added over time, and as such not all questions had responses from all students. (b) Pie chart of class standing when the class was taken (n = 107). (c) Bar plot of students’ prior coding experience by programming language (n = 83). (d) Pie chart of students’ prior research lab experience (n=42).

Out of 83 students, 72 of the 83 students had some familiarity with at least one programming language (Fig 1c). The most common language was C++, likely due to the computer science course requirements for the QBIO major. Only 28.9% of students had experience with R and 43.4% of students had experience with Python. Although 86.7% of students had some level of programming knowledge, many mentioned that they lacked experience combining programming with biological data analysis. One student stated that, “of the courses I’ve taken at USC so far, I’ve taken biology courses and I’ve taken computer science courses, but I really want to take a course that involves both so I can better understand how the subjects work together in a computational biology research lab setting.”

Of 42 students asked across two semesters, 21 did not have a current research lab position when they took the pre-semester survey (Fig 1d). Many students mentioned that they wanted to take MODA and gain skills to join a research lab. One student said that “[they hoped] to gain a strong foundation in using R and Python for data analysis, which will be especially beneficial to transitioning to research.” To facilitate this transition, part of the MODA curriculum focused on guiding students through finding relevant labs and contacting potential PIs. Instructional staff provided advice on writing emails and resumes to interested students.

In the pre-semester survey, students were also asked what skills they were most interested in learning (Table 1). These skills broadly fell into four categories: computational skills, research experience, scientific communication, and community building. These responses were used to inform minor year-to-year curriculum changes for the class. Even though 86.7% of students had some prior exposure to coding languages, 42.2% of students still indicated that learning computational skills was a strong driver of their participation in MODA.

**Table 1.** Student responses to a pre-semester survey indicating desired outcomes for MODA (n=83).

| Category | Specific outcomes | % student responses |
| --- | --- | --- |
| Computational skills | General knowledge | 42.2 |
|  | Learn R | 27.7 |
|  | Learn Python | 22.9 |
|  | Machine learning skills | 4.82 |
|  | Learn GitHub | 3.61 |
| Research experience | General experience | 51.8 |
|  | Application of skills to real biological data | 25.3 |
|  | Find research opportunities | 10.8 |
|  | Collaborative work | 3.61 |
| Scientific communication | Reading and writing scientific papers | 6.02 |
|  | Presenting research | 1.2 |
| Community building | Explore bioinformatics field | 13.3 |
|  | Get to know other students in degree program | 10.8 |

### Multi-Omic Data Analysis successfully improves student self-assessment of competence in bioinformatics skills

We first aimed to establish the effectiveness of our PTP-based class as an introductory-level bioinformatics course. Students were asked to assess their personal competence with the skills taught in MODA in the post-semester survey. A larger proportion of students said they felt proficient in computational and research skills across the board after taking the class (Fig 2a). This improvement is higher in coding skills, as we see an increase in the number of students who reported comfort with basic R from 40.4% before the course to 100% after. In a similar manner, students’ comfort with basic Python increased from 47.6% to 78.6% after the course was taken. More relevant to bioinformatics-specific skills are the number of students who reported comfort with “Scientific R” and “Scientific Python”, which refer to the use of packages and libraries that are commonly utilized in bioinformatics analysis (e.g. NumPy, SciPy, scikit-learn, etc. in Python and DESeq2, SummarizedExperiment, maftools, etc. in R). Less than 20% of students in both categories stated that, prior to the course, they would have been proficient with such skills. The large difference between the number of students with self-reported proficiency in basic R/Python and those with self-reported proficiency in scientific R/Python is indicative of the fact that while students within bioinformatics programs like QBIO may have some exposure to coding languages, this experience is void of any practical applications of such knowledge as it relates to bioinformatics research.

**Fig 2.**
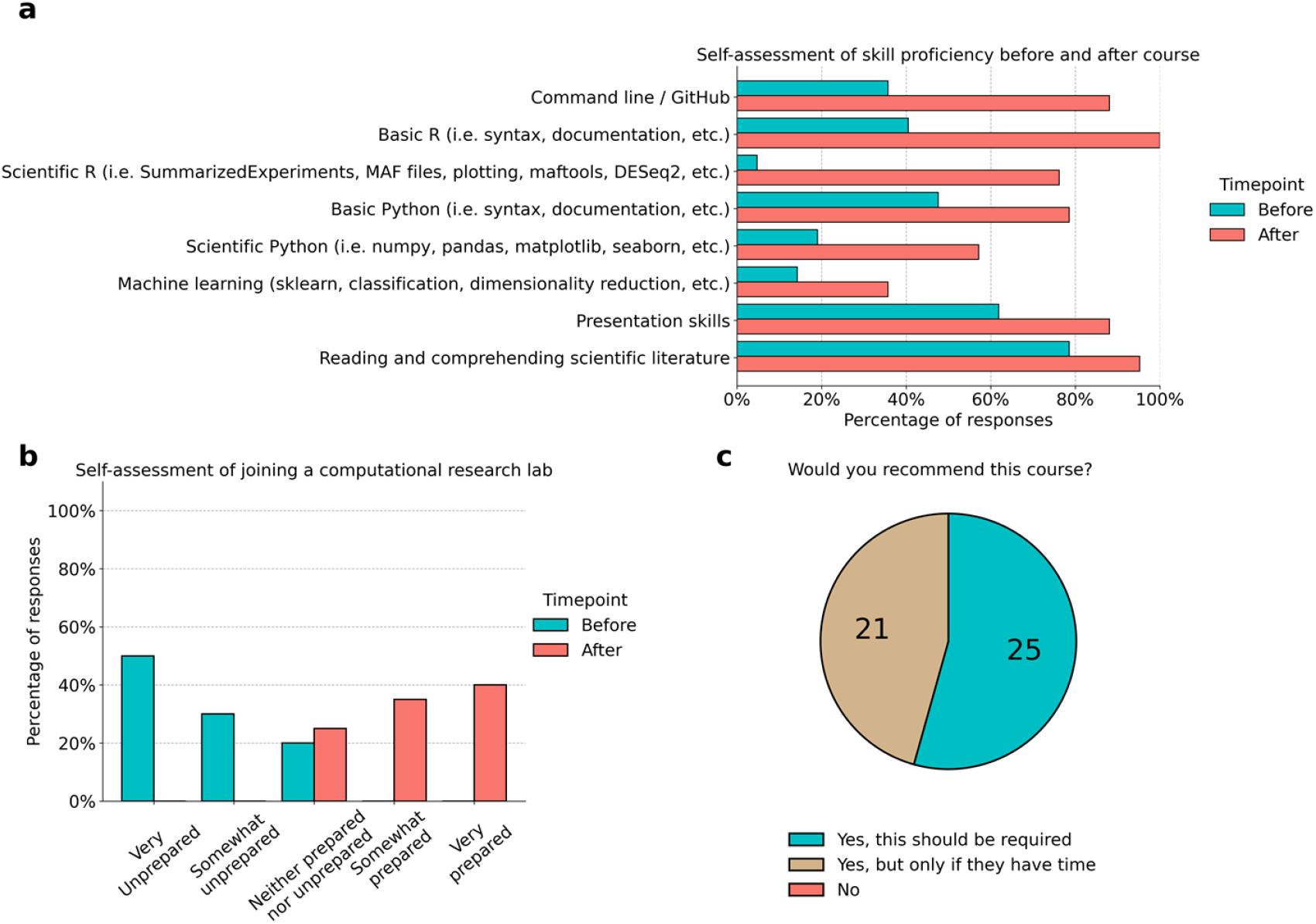
Students post-MODA report improved bioinformatic skills and confidence in research ability. (a) Student assessed their proficiency in eight topics covered by MODA, both before and after participation in the course (n=42). For each topic at each timepoint, students indicated if they were “Comfortable With” or “Not Comfortable With” said topic. (b) Students reported a self-assessment of their perceived preparedness for joining a computational research lab as an undergraduate student (n = 20). (c) Students were asked if they would recommend MODA to other students (n = 46). Students had the option to qualify a positive response with whether they believe it should be required of undergraduate QBIO students.

This interpretation is further supported by the fact that students reported a greatly improved self-assessment of their preparedness to join a computational research lab as an undergraduate student. No student rated themselves lower than “Neither Prepared nor Unprepared” for joining a research lab after taking MODA, whereas every student rated themselves at or lower than “Neither Prepared nor Unprepared” when considering their skill level prior to the course (Fig 2b). We also see a clear skew left in “After” course ratings, where most students considered themselves “Somewhat Prepared” or “Very Prepared” to contact professors for potential computational research positions. On an individual level, every student rated themselves at higher preparedness after taking the course. Most students also chose to explain their rating, with one student stating that “[if] not for this class, I would definitely feel more imposter syndrome, and my lack of confidence likely would have prevented me from applying to labs.” Multiple other students state similar perspectives (see data availability). Even among students who were in research labs prior to taking MODA, the vast majority still felt that the class was useful for both their work in lab and their overall bioinformatics skill level. One student said, “I liked being exposed to a broader range of computational methods, compared to previously being mostly focused on what I was doing in the lab.” One student additionally stated that they “would have liked to take this course before beginning in my lab”, a sentiment that was echoed by multiple others. Overall, MODA appears to successfully encourage confidence with bioinformatics skills and computational research.

### PTP positively affects student engagement and comfort with learning bioinformatics skills

In the post-semester survey, students were asked what they believed to be the most valuable aspects of MODA. While the most common responses highlighted hard skills such as coding (41.5%) and research experience (32.1%), many students mentioned that they enjoyed the community building aspects of the course (Table 2). One student said of the course, “I’ve never seen the level of [camaraderie] I saw in this class in ANY other class I’ve taken. I think also learning from my peers aside from the instructors was a big advantage. And also, the instructors being around our age… made it easier to ask for help or simply admit that I didn’t understand something.” This supports the case that PTP learning strategies benefit from interpersonal relationships that can form when students and student instructors are on the same educational level. Another student mentioned “I think the student aspect of the group has been super helpful, I’m very comfortable with asking for help. I’ve also learned so much in this group that I can actually apply to my career in the future.” Students feeling more comfortable asking for help demonstrates how PTP learning can improve student learning by improving perceived accessibility of educational assistance.

**Table 2.**
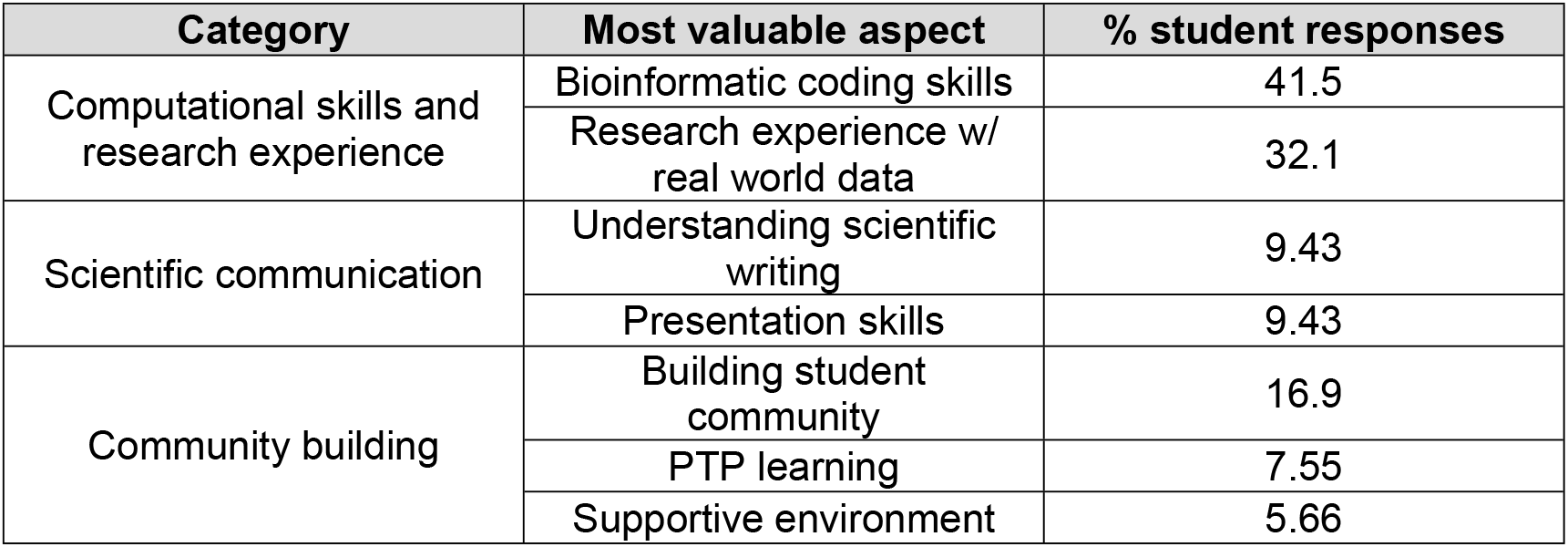
Post-semester survey responses on the most valuable aspect of MODA participation (n=70).

| Category | Most valuable aspect | % student responses |
| --- | --- | --- |
| Computational skills and research experience | Bioinformatic coding skills | 41.5 |
|  | Research experience w/ real world data | 32.1 |
| Scientific communication | Understanding scientific writing | 9.43 |
|  | Presentation skills | 9.43 |
| Community building | Building student community | 16.9 |
|  | PTP learning | 7.55 |
|  | Supportive environment | 5.66 |

This open attitude is well reflected in students’ responses to whether they would recommend this course to other students. Out of the 30 students who responded, every single student responded that they would recommend the course (Fig 2c). Slightly above half of the students said that the course should be a requirement of the degree program.

### Course participation affects long-term career goals in bioinformatics

MODA also appeared to have an impact on students even months to years after participation. Most alumni who responded to the alumni survey reported continued use of the skills and knowledge they learned in MODA, with the most used skills being “Scientific Communication”, “R”, and “Python” (Fig 3a). More significantly, however, were the reported impacts of MODA on personal career goals and overall perception of bioinformatics as a field. In the alumni survey, nearly half of respondents indicated that MODA increased their interest in pursuing bioinformatics post-undergrad, with about a third stating that MODA exposed them to careers related to bioinformatics (Table 3).

**Table 3.** Alumni survey responses on the impact of MODA participation (n=13).

| Impact of MODA | Outcome | % alumni responses |
| --- | --- | --- |
| Positive | Interest in pursuing bioinformatics | 46.2 |
|  | Improved perception of bioinformatics | 30.8 |
|  | Exposure to careers related to bioinformatics | 30.8 |
|  | Interest in data analysis | 23.1 |
|  | Improved ability to find research opportunities | 15.4 |
|  | Pursuing a computational biology Ph.D. | 7.69 |
| Negative | Prefer medicine over research | 38.5 |
|  | Prefer C.S. over biology | 7.69 |
|  | No interest in data analysis | 7.69 |

**Fig 3.**
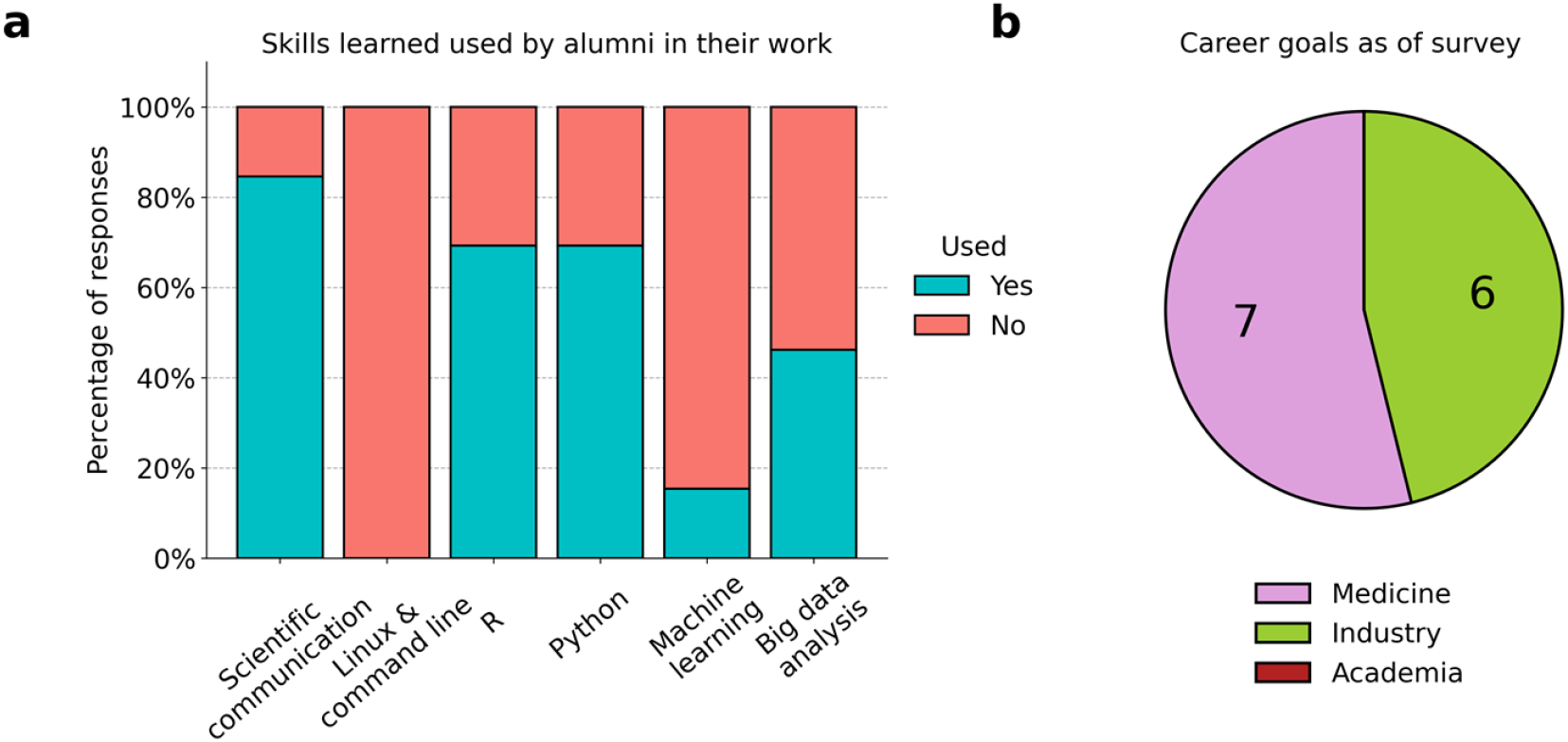
Alumni responses to questions about their overall experience in MODA. (a) Alumni indicated which skills taught in MODA that they continue to use past participation in the course (n = 13). (b) Pie chart of responses for which area best described their career goals at the time of taking the survey. No responses indicated “Academia”.

Almost a third of responses also indicated an improved perception of bioinformatics, and a quarter indicated that participation in the course confirmed or created an interest in data analysis. Around 38.5% of respondents did state that MODA made them realize they preferred medical practice over research, and we do see that about half of respondents indicated that they currently had or aimed to have a career in medicine (Fig 3b). One of the 13 alumni said that they “realized that [they] didn’t like the actual implementation of big data analysis” as much as they thought they did. Another respondent realized they preferred the coding work in MODA so heavily that they switched their major to Computer Science. On the other hand, one of the respondents also indicated that their experiences with research in MODA led to their current position as a Ph.D. student, where they continue to work with multi-omic data. Even if we don’t consider whether MODA had a “positive” or “negative” effect on students’ perceptions of and participation in bioinformatics, it is evident that MODA has played a significant role in the education and career development of many students. Half of the respondents still indicate a continued interest in bioinformatics work, with that interest leading to careers in the private sector (Table 3 and Fig 3b).

To investigate the long-term attitudes toward PTP learning, alumni survey respondents were asked their opinion on the peer-led aspect of the course. 46.2% of respondents indicated that it made content more approachable and improved communication between students and student instructors (Table 4). Two students mentioned that the lack of student instructor experience was apparent, but most responses were overwhelmingly positive. This provides support to the idea that even though student instructors may have less experience in teaching, the interpersonal communication and community building benefits of PTP learning may outweigh any disadvantages by lowering stakes and improving learning outcomes.

**Table 4.**
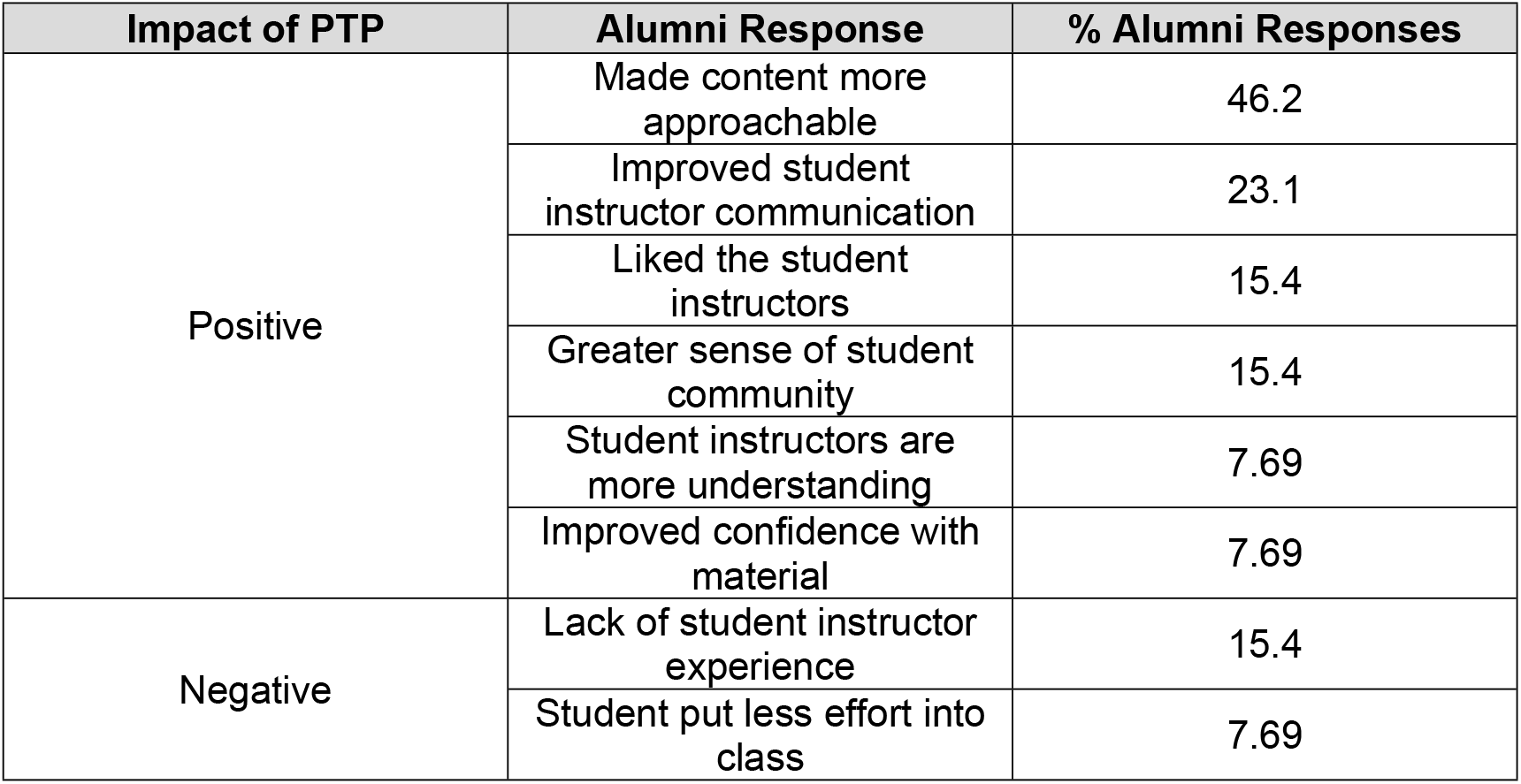
Alumni responses to the impact of PTP on their MODA experience (n = 13).

| Impact of PTP | Alumni Response | % Alumni Responses |
| --- | --- | --- |
| Positive | Made content more approachable | 46.2 |
|  | Improved student instructor communication | 23.1 |
|  | Liked the student instructors | 15.4 |
|  | Greater sense of student community | 15.4 |
|  | Student instructors are more understanding | 7.69 |
|  | Improved confidence with material | 7.69 |
| Negative | Lack of student instructor experience | 15.4 |
|  | Student put less effort into class | 7.69 |

## Discussion

In this case study, we evaluated the objectives and execution of a PTP-learning introductory bioinformatics course in the QBIO undergraduate degree program. We found that the student-led course, MODA, was anecdotally effective at addressing barriers to pursuing computational biology research as an undergraduate student. Students left the course with improved self-assessment of bioinformatics skills across multiple areas of focus (R, python, scientific communication, etc.) as well as a greater understanding and appreciation for the role of computational work in biological and biomedical sciences. The PTP-learning aspects of MODA were both directly and indirectly cited by multiple students as having a significant positive impact on their learning and performance in the course. Overall, we argue that MODA successfully accomplished the course’s primary goals of improving student confidence in bioinformatics work, encouraging the value of computational work in biology, encouraging students to participate in computational biology research labs, and building a community among students via the application of peer-led learning.

MODA addresses the need for approachable and beginner-friendly bioinformatics courses that can prepare students for work in a computational biology research laboratory, which we argue has been missing from large parts of bioinformatics higher education. Much of the content in MODA receives little to no coverage in traditional biology classes. While practical computational skills may understandably be outside the purview of a general biology course, this only serves to further highlight the importance of courses like MODA. Even courses specifically created for the QBIO major at USC tend to provide students with limited practical experience, not to mention are predominantly taken by students in their third or fourth year of college. Examples of concepts in MODA that have limited practical coverage in other USC computational biology courses include Kaplan-Meier plots, differential expression analysis, multi-omic data and analysis, Bash scripting, GitHub, popular R and Python packages in bioinformatics, etc.

We can attribute the gaps in bioinformatics education to being reflective of a larger systemic issue in academic biological research. There is a well-documented cultural and professional divide between traditional benchwork scientists and computational scientists, with the most memorable manifestation of this tension being the concept of “research parasites” introduced in the infamous 2016 New England Journal of Medicine editorial “Data Sharing” [12]. This divide typically sidelines bioinformaticians and computational biologists in more traditional “wet” labs, largely devaluing their contributions and expertise [13]. The interdisciplinary nature of bioinformatics and computational biology challenges the mores of biological research, and thus we see a similar divide in the education of aspiring scientists.

MODA utilizes PTP-learning to address this divide at a grassroots level, where more experienced students draw on their own skills and understanding of bioinformatics to help prepare newer students for computational biology research work. Students have the benefit of greater freedom to explore interdisciplinary fields and concepts due to the nature of the collegiate experience, and as such can bring a more collaborative and open-minded energy to teaching and learning. Having multiple students on the instruction staff at a time also means students with different types of research experience, both “wet” and “dry”, contribute to the course. The result is a constantly evolving but comprehensive and beginner-friendly curriculum that instructors determine by merely asking themselves, “What do I wish I knew before joining a lab?”. As bioinformatics and computational biology are rapidly developing fields, the timely experiences that students bring to the table mean the MODA curriculum can expose newer students to the hard and soft skills most relevant to current research work at the undergraduate level.

While we have successfully established MODA in the QBIO undergraduate degree program, this project only serves as a case study. This specific application of PTP bioinformatics education benefits from certain advantages due to our host institution, such as a strong research environment, supportive undergraduate department, and access to student instructors with prior bioinformatics research experiences. Whether or not other institutions have these advantages will impact the feasibility of starting a program like MODA. PTP-learning itself also has some potential limitations. For one, PTP-learning structures by nature have fast student instructor turnover, where student instructors can theoretically change every semester. This means PTP-learning programs like MODA are especially sensitive to waning student engagement, and semester-to-semester momentum is vital to the longevity of the course. Students also cannot be full-time employees of the institution they attend, so pay can be a barrier to becoming a student instructor, as the role essentially amounts to a part-time job. While solutions to these challenges would likely vary from institution to institution, our implementation of MODA included the following: The course was initially a student club that began online in Spring 2021 during the COVID-19 quarantine, and as such, student pay was not considered. When the program became available as an official course in Spring 2022, student instructors were paid as student employees by the department. At the time of this paper, we have not struggled to find students interested in teaching each semester. Potential incoming students are heavily recruited through email before and at the start of each semester and taking the course for research credit is a significant incentive.

Regardless of the limitations to PTP-learning, we still find in this case study that the benefits of PTP-learning in introductory bioinformatics education are worth the effort. To expand the impact of MODA and further evaluate its effectiveness, we ideally would make the course available to students in traditional biology and computer science degree programs, and not just students in QBIO. This would be one advantage of PTP-learning programs remaining student organizations, as clubs would allow more flexibility in which students could participate. This would also further encourage interdisciplinary discussion and encourage positive perceptions of computation in biological research.

In conclusion, MODA addresses a current gap in bioinformatics education with PTP-learning. While MODA is inherently limited in its generalizability to other students and other institutions as a case study, the program does demonstrate the potential for peer-led teaching to fill an educational gap and improve interdisciplinary understanding between traditionally and culturally isolated fields of study.

## Methods

### Multi-omic data analysis course structure

Multi-Omic Data Analysis (MODA) is a student-led cancer bioinformatics course hosted by the USC Department of Quantitative and Computational Biology (QCB) and run by students in the Quantitative Biology (QBIO) undergraduate degree program. The goal of the course is to provide QBIO undergraduate students with the foundational skills and experience necessary for research and careers in computational biology and bioinformatics, as well as build a greater sense of community within the degree program. The program began as a student club during the Spring 2021 semester and was hosted entirely online due to the COVID-19 pandemic. Starting in Fall 2021, course lectures were held in-person at USC. Beginning in Spring 2022, MODA was offered with the option of taking the class for research credit, which allowed students to complete 2 of the 6 research units required for graduation.

The course is structured like a standard college-level course. Schedules are altered each semester to ensure the maximum number of interested students can participate but generally include two 1-hour lectures each week (sample course calendar available in supporting information). An additional 3-5 hours of office hours are also offered, depending on the number of instructional staff per semester.

All student instructional staff are current QBIO students who previously participated in the course and are selected via an application process by current student instructional staff. Student instructional staff include one to two student instructors, who are responsible for course lectures, assignments, and projects. Two to three student teaching assistants are also selected and are responsible for assignment grading, review sessions, and guest panels. Finally, the student course director position is held by the most senior member of the student instructional staff, and is responsible for logistical operations, such as scheduling, communicating with faculty and guest speakers, and recruiting prospective students to the course.

MODA has four main objectives for participating students: foundational knowledge of bioinformatics and multi-omic research (e.g. multi-omic data types, command line/Linux, accessing publicly available data from The Cancer Genome Atlas, etc.), developing skills in R, developing skills in Python, and experience in scientific communication (sample course description available in supporting information). Students engage with these four objectives through lectures and weekly assignments, as well as a final project on a bioinformatic cancer research question of their choice The final project serves as a culmination of all the skills covered in MODA, and as an opportunity for students to demonstrate their ability to structure a small research project and present their findings. At the end of the semester, final project groups (3-4 students) give a 15–20-minute presentation of their project to student instructional staff, their fellow students, and a panel of faculty graders. The presentations and post-event mixer simultaneously function as a community event for MODA, the QBIO major, and the QCB department.

While this course is graded for credit, attendance accounts for 40% of the grade, with the other 60% including assignments, a literature review presentation, an R review project, a Python review project, and the final project presentation and paper. Assignments are graded on completion and effort, rather than accuracy, to reduce stress on students. If the students enrolled in the course for research credit, their course final grade is included in their official transcript. All coding assignments and projects are completed by students using their personal laptops. Other resource requirements for MODA are access to the internet to download data from The Cancer Genome Atlas and Clinical Proteomic Tumor Analysis Consortium, installation of R/RStudio and Python, and a Linux terminal.

### Survey data collection and analysis

Before the start of each semester, 83 incoming students completed pre-semester surveys to collect information on the prior experience and goals of those interested in MODA. A pre-semester survey was not offered to the inaugural cohort in Spring 2021.

At the end of each semester, students completed post-semester surveys to collect information on students’ self-evaluation of their skills after participation in the course, as well as collect feedback on their overall experience. This survey was not implemented in Fall 2022. A total of 70 student responses were collected. Unlike the pre-semester survey, the post-semester survey experienced minor alterations for each semester, primarily for the inclusion of more questions related to student self-evaluation. Many survey questions were also optional. As such, most questions have a sample size of student responses smaller than 70. For both the pre-semester and post-semester surveys, students were de-identified.

The alumni survey was offered to students who participated in MODA between Spring 2021 and Spring 2023, with the goal of collecting information on the post-course impact of MODA, as well as specific inquiries on the impact of the PTP aspect of MODA. A total of 13 anonymous responses were collected. A full list of questions used in all surveys can be found in the supporting information.

Categorical responses (i.g. “Before taking this class, were you in a lab?”, “How prepared do you feel about joining a lab after this class?”, etc.) were plotted as bar plots and pie charts. One-hot encoding was used for analysis purposes if necessary. Plots were made using Python 3.11.4 and packages pandas 1.5.3 and NumPy 1.26.4 for data processing. All bar plots and pie charts were created using matplotlib 3.8.0. Sankey plots were made using sankeyflow 0.3.8.

Most responses were text responses; as such, they were processed and analyzed manually by grouping responses by content. Text responses were then plotted in text tables using grid 4.3.1 and gridExtra 2.3 in R 4.3.1 and RStudio 2023.6.1.524.

## Acknowledgements

We would like to thank all the students, past and present, who continue to participate and contribute to the MODA program, for whom this program is for and could never exist without. Thank you to our former and current instructing staff: David Wen, Haley Boren, Echo Tang, Jonathan Martinez, Olivia White, Brandon Ye, Erika Li, Akansha Sallakonda, Jeanne Revilla, and Eros Mendoza. Special thanks to program founders Kate Manley and Terrence Li, for developing and executing the concept of this course. Thank you to the Ellison Medical Institute LLC and especially Dr. Jerry Lee for their continued support and use of their facilities. Finally, we’d like to thank Dr. Remo Rohs, Dr. Peter Calabrese, and the rest of the USC Department of Quantitative and Computational Biology faculty members for their support and participation in this program.

## Supporting information

**S1 File. Example course calendar**

**S2 File. Course description**

**S3 File. Survey questions**

## References

1. Precedence Research. Bioinformatics Market Size to Hit Around US$45.6 Bn by 2030 Global Newswire: Global Newswire; 2022 [cited 2024 Aug 4].

2. Way GP, Greene CS, Carninci P, Carvalho BS, de Hoon M, Finley SD, et al. A field guide to cultivating computational biology. PLOS Biology. 2021; 19(10):e3001419. doi: 10.1371/journal.pbio.3001419.

3. Integrated Postsecondary Education Data System. In: Statistics NCfE, editor. nces.ed.gov2022.

4. Perkins KK, Adams WK, Pollock SJ, Finkelstein ND, Wieman CE. Correlating Student Beliefs With Student Learning Using The Colorado Learning Attitudes about Science Survey. 2004 Physics Education Research Conference. 2004; 790:61–4.

5. Knopes J, Cascio MA, Warner B. Intraprofessionalism and Peer-to-Peer Learning in American Medical Education. Qual Health Res. 2024; 34(6):528–39. Epub 20231211. doi: 10.1177/10497323231218137. PubMed PMID: 38079522.

6. Surabenjawong U, Phrampus PE, Lutz J, Farkas D, Gopalakrishna A, Monsomboon A, et al. Comparison of Innovative Peer-to-Peer Education and Standard Instruction on Airway Management Skill Training. Clinical Simulation in Nursing. 2020; 47:16–24. doi: 10.1016/j.ecns.2020.06.009.

7. Maccabe R, Fonseca TD. ‘Lightbulb’ moments in higher education: peer-to-peer support in engineering education. Mentoring & Tutoring: Partnership in Learning. 2021; 29(4):453–70. doi: 10.1080/13611267.2021.1952393.

8. Hannah C, James S, Elizabeth W. Undergraduate research as a capstone requirement. Involve: A Journal of Mathematics. 2014; 7(3):273–9. doi: 10.2140/involve.2014.7.273.

9. Canaria JA, Schoffstall AM, Weiss DJ, Henry RM, Braun-Sand SB. A Model for an Introductory Undergraduate Research Experience. Journal of Chemical Education. 2012; 89(11):1371–7. doi: 10.1021/ed200582v.

10. Maxwell MC, Snyder JJ, Dunk RDP, Sloane JD, Cannon I, Wiles JR. Peer-led team learning in an undergraduate biology course: Impacts on recruitment, retention, and imposter phenomenon. BMC Research Notes. 2023; 16(1):73. doi: 10.1186/s13104-023-06338-7.

11. The Cancer Genome Atlas Research Network, Weinstein J, Collisson EA, Mills GB, Mills Shaw KR, Ozenberger BA, et al. The Cancer Genome Atlas Pan-Cancer analysis project. Nature Genetics; 2013. p. 1113–20.

12. Longo Dan L, Drazen Jeffrey M. Data Sharing. New England Journal of Medicine. 2016; 374(3):276–7. doi: 10.1056/NEJMe1516564.

13. Lewis J, Bartlett A, Atkinson P. Hidden in the Middle: Culture, Value and Reward in Bioinformatics. Minerva. 2016; 54(4):471–90. Epub 20160711. doi: 10.1007/s11024-016-9304-y. PubMed PMID: 27942075; PubMed Central PMCID: PMCPMC5124041.

